# Disadvantages of writing, reading, publishing and presenting scientific papers caused by the dominance of the English language in science: The case of Colombian PhD in biological sciences

**DOI:** 10.1101/2020.02.15.949982

**Authors:** Valeria Ramírez-Castañeda

**Affiliations:** Department of Philology, University of Barcelona, Barcelona, Spain; Department of Integrative Biology, University of California, Berkeley, United States of America; Museum of Vertebrate Zoology, University of California, Berkeley, United States of America

## Abstract

The success of a scientist depends on their production of scientific papers and the impact factor of the journal in which they publish. Because most major scientific journals are published in English, success is related to publishing in this language. Currently, 98% of publications in science are written in English, including researchers from English as a Foreign Language (EFL) countries. Colombia is among the countries with the lowest English proficiency in the world. Thus, understanding the disadvantages that Colombians face in publishing is crucial to reducing global inequality in science. This paper quantifies the disadvantages that result from the language hegemony in scientific publishing by examining the additional costs that communicating in English creates in the production of articles. It was identified that more than 90% of the scientific articles published by Colombian researchers are in English, and that publishing in a second language creates additional financial costs to Colombian doctoral students and results in problems with reading comprehension, writing ease and time, and anxiety. Rejection or revision of their articles because of the English grammar was reported by 43.5% of the doctoral students, and 33% elected not to attend international conferences and meetings due to the mandatory use of English in oral presentations. Finally, among the translation/editing services reviewed, the cost per article is between one-quarter and one-half of a doctoral monthly salary in Colombia. Of particular note, we identified a positive correlation between English proficiency and higher socioeconomic origin of the researcher. Overall, this study exhibits the negative consequences of hegemony of English that preserves the global gap in science. Although having a common language is important for science communication, generating multilinguistic alternatives would promote diversity while conserving a communication channel. Such an effort should come from different actors and should not fall solely on EFL researchers.

## Introduction

At the same time that scientific articles became the measure of scientific productivity, English was imposed as the language of science, culture, and the global economy (Johnson et al., 2018). As a consequence, today 98% of publications in science are written in English, especially in the areas of natural and basic sciences, establishing English as the *lingua franca* of science (Gordin, 2015). This creates a disadvantage for scientists with English as a Foreign Language (EFL) because they must publish complex texts in a foreign language to advance their careers (Parchomovsky, 1999). This disadvantage gives rise to global inequalities, especially in countries where the majority of the population receives minimal English training and bilingualism with English is very low (Tardy, 2004). Thus, English proficiency and socioeconomic level influence scientific success, access to knowledge and expatriation, among others.

One of the most important goals for modern society is to increase scientific production from Africa, Latin America, Middle East, and developing Asia. There is a strong correlation among English proficiency, economic development, and technological innovation in terms of number of articles, number of researches and research and development expenditure (EF Education, 2018). Therefore, the prevalence of the English language in the sciences deepens the inequality in knowledge production between countries with high and low English proficiency (Dei & Kempf, 2006), maintaining the gap in scientific production between the countries of the global south or peripheral and the countries of the global north (include the G8 countries and Australia), reducing the individual scientific contributions of EFL scientists (Murphy & Zhu, 2012). Together these factors limit the advancement of the broad scientific communities within those countries (Whatmore, 2009).

Numerous studies have identified the use of English in academia as a source of inequality and segregation in science (Flowerdew, 1999; Pabón-Escobar & da Costa, 2006; Muresan & Pérez-Llantada, 2014; Curry & Lilis, 2017, Hanauer et al., 2019). These inequities affect the scientific community at multiple levels. In local communities of EFL countries, scientific thinking is harmed, particularly in higher education, as learning depends on cultural attitudes derived from the native language spoken by the students, and science becomes alien to their own experiences (Gulbrandsen et al., 2002; Lee & Fradd, 2007; de Vasconcelos, 2006). Diversity in language promotes diversity in thinking, affecting creative process and imagination; thus, the maintenance of multilingualism in science could have an impact on scientific knowledge in itself (Lee & Fradd, 2007).

Local journals are a refuge for communication of scientific research in languages other than English, nevertheless they are often perceived as low-quality, since the most important research work is often reserved for international journals. Therefore, readers with language barriers only have access to this kind of knowledge and are even unaware of the most significant research being conducted in their own region, which has resulted in a void in information important for political decision making, regional environmental policies and conservation strategies, within the territories and a bias for global decision making (Salager-Meyes, 2006; Alves & Pozzebonm, 2014; Amano et al., 2016). In addition, despite the importance of local knowledge, the professional success of a scientist correlates to a greater extent with their “internationalization”. This constant pressure could be influencing academic migration, known as “expatriation”. English learning is one of the pressure factors of migration, as it is more difficult to achieve upper English proficiency for scientists who remain in EFL countries (Benfield & Howard, 2000; Ferguson, 1999; de Vasconcelos, 2006).

In periphery countries there is a strong relationship between English proficiency and socioeconomic origin, thus it is important to understand the publishing costs associated with the socioeconomic origin of the doctoral student. Among Latin America, Colombia is the second most unequal territory: in 2018 it invested only 0.24 % of its GDP (Sweden investment was 2.74% of its GDP) in science, technology and innovation (Portafolio, 2019), and it has one of the lowest levels of English proficiency among the world rankings (EF Education, 2018). In addition, only 0.015% of the population has or is doing a doctorate in basic or natural sciences (UNESCO, 2016). This study aims to determine if Colombian doctoral students of natural sciences face disadvantages when publishing scientific articles in English, compared to publications in their first language, and to quantify the extra work that these scientists put into writing, reading, and presenting their work in English. In addition, this study examines the impact of socioeconomic background on English proficiency and the costs it generates when publishing.

## Materials and Methods

In order to determine the costs of publishing in English experienced by Colombian doctoral students and doctorates in biological sciences, 49 doctoral students participated in the “Implications of language in scientific publications” survey containing 44 questions in Spanish language (S3 Survey questions). Additionally, the prices offered by prestigious scientific publishers for translation (Spanish to English) and editing of scientific texts were searched to measure the economic impact in relation to a Ph.D. student salary in Colombia.

### Survey Construction

The main survey of this work, entitled “Implications of language in scientific publications,” has 44 questions divided into three sections: basic data, writing articles in English, and learning English (S3 Survey questions). The responses obtained were grouped for statistical analysis (Table 1).

**Table 1.** Summary of survey questions divided by groups used for correspondent statistical analysis.

| Group | Variable | Type of variable | Reference |
| --- | --- | --- | --- |
| **Socioeconomic data** | Differentiation between attending public and private education in the high school and university | Binary (public/private) | Cataño, 2017 |
|  | Occupation of the father and mother to define socioeconomic status | Categorical | Requena, 2012 |
|  | Socioeconomic stratification in Colombia from him- or herself, father, and mother | Ordinal (1 to 6) | The Congress of Colombia, 1994 |
| **English proficiency** | First language(s) | Categorical |  |
|  | Language of the country in which they live | Categorical |  |
|  | Proficiency in writing, oral expression, reading and listening in English | Ordinal (bad, regular, good, excellent) |  |
|  | Age of beginning of English learning | Discrete quantitative |  |
|  | Years studying English | Discrete quantitative |  |
|  | Satisfactory experience in the English learning institution | Ordinal (1 to 5) |  |
|  | Years lived in English-speaking country | Discrete quantitative |  |
|  | Percentage of English speaking daily | Percentage quantitative |  |
| **Publications** | Number of scientific publications reviewed by academic peers | Discrete quantitative |  |
|  | Percentage of those publications in English or in a language other than English | Percentage quantitative |  |
| **Costs of writing in English** | Payment or favors for editing or translating scientific publications | Binary (yes/no) |  |
|  | Percentage of articles - payment or favors for editing or translating scientific publications | Percentage quantitative |  |
|  | Number of rejections in scientific journals associated to English writing | Discrete quantitative |  |
|  | Time spent writing a scientific article in English or Spanish (labor hours) | Continuous quantitative |  |
|  | Preference between writing directly in English or translating to English | Binary (English/translating) |  |
| **Writing difficulties in English or Spanish** | Difficulty of writing scientific paper introduction, methods, results, discussion, conclusion | Ordinal (1 to 5) |  |
| **Reading difficulties in English or Spanish** | Difficulty in understanding general text / Understanding scientific terminology / Interpreting figures and tables | Ordinal (1 to 5) |  |
| **Participation in conferences** | Oral presentations in international conferences | Binary (yes/no) |  |
|  | Level of anxiety in oral presentations in English or Spanish | Ordinal (1 to 5) |  |

### Statistical analysis

Statistical analyses were performed in R v.3.6.1 (R Development Core Team, 2011) and data were plotted with the ggplot package (Valero-Mora, 2015). To compare reading and writing between English and Spanish, time investment and the level of anxiety in conferences participation, an ANOVA was performed (*aov* in package ‘stats’ v3.5.3). The margin of error was calculated with 95% confidence. An Analysis of Principal Components (PCA) was performed using the variables contained in the “English proficiency” and “Socioeconomic data” groups for reducing redundancy in the variables (*PCA* in package FactorMiner v2.2). The proportion of variance explained by each principal component was reviewed, and only the first principal component was retained for each dataset, as it described 51% and 62% correspondingly of the total variation. Subsequently, a linear regression was executed with the intention of comparing these two variables, English proficiency PC1 vs socioeconomic status PC1 (using *lm* in package ‘stats’ v3.5.3).

### Editing and translation services costs

In order to visualize the prices of English editing and translation services for scientific texts, information was sought in five of the most relevant scientific publishers (SAGE language services, 2018; Cambridge University Press Author Services, 2018; Elsevier author services, 2018; SpringerNature author services, 2018 and Wiley editing services, 2018). The information and costs of these services are public and can be obtained through the web pages of publishers. All data were taken with respect to prices for a text of 3000 words, as that is the average length of a scientific article; searches were performed in October 2018.

These publishers offer two types of editing services, a three-day service (Premium) and a one-week service (standard); both prices were used for the analysis. Only the prices for Spanish - English translations were used. Finally, these prices were compared with an average doctoral salary in Colombia (Colciencias, 2019), 947 US dollars or 3 million Colombian pesos (1 US dollar = 3.166 Colombian pesos, exchange price on January 31, 2019).

## Results

A total of 49 responses were obtained from Colombian doctoral students or doctorates in biological sciences whose first language is Spanish. From Colombians’ surveyed 92% (sd = 0.272) of their published scientific articles are in English and only 4% (sd = 0.2) of their publications were in Spanish or Portuguese. In addition, 43.5% of the doctoral students stated at least one rejection or revision of their articles because of the English grammar.

With regards to time investment, there was a significant increase in the time invested writing a scientific article in English in comparison to Spanish for survey participants (Fig 1). The process of writing in Spanish takes on average 114.57 (sd = 87.77) labor hours, while in English, 211.4 (sd = 182.6) labor hours. On average, these scientists spend 96.86 labor hours more writing in English. However, 81.2% of the doctoral students stated that they prefer to write directly in English in comparison to writing in Spanish and then translating into English.

**Fig 1.**
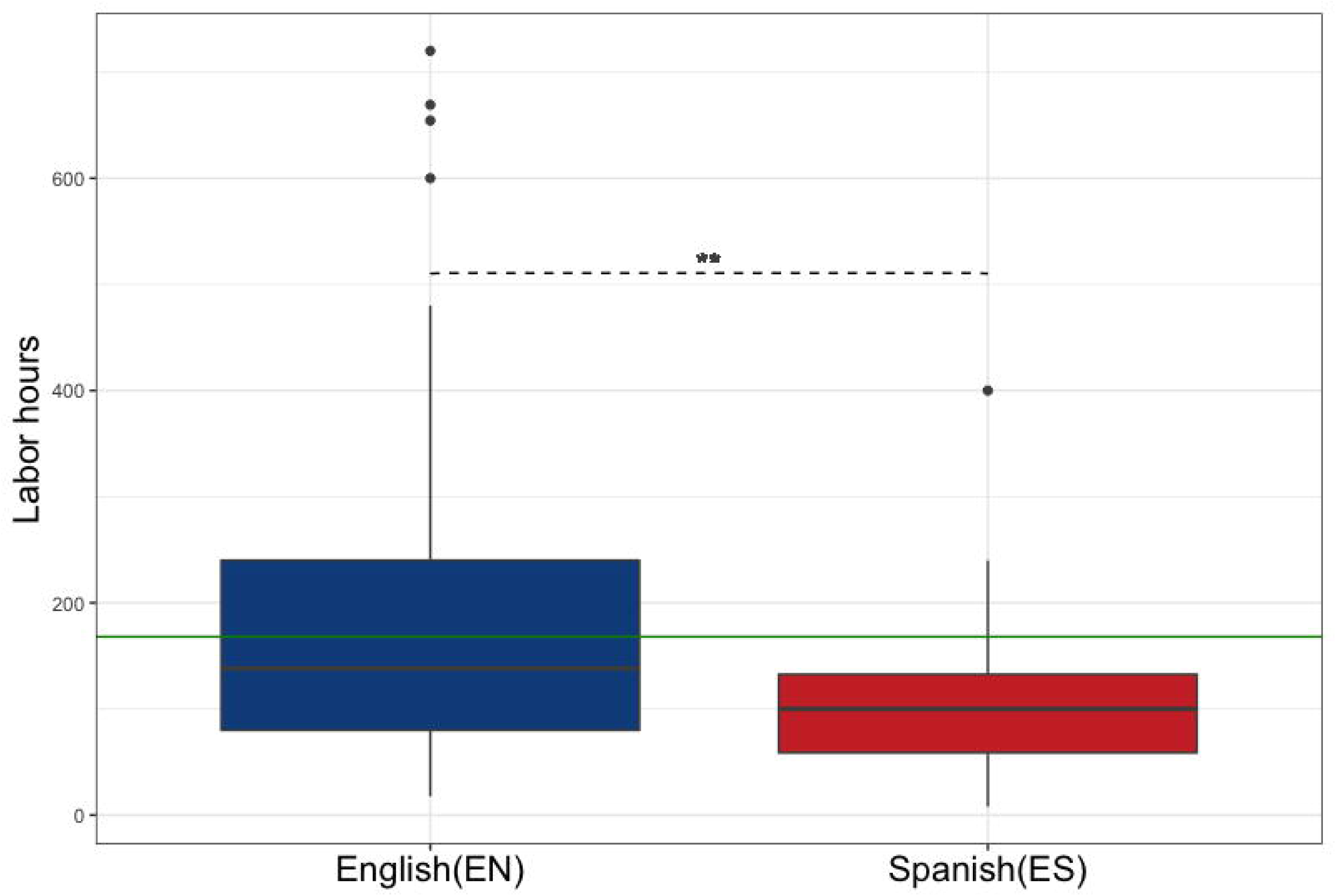
Time invested writing a scientific article in English (EN) as a secondary language and in Spanish (ES) as a first language. An ANOVA analysis was performed to compare the variables obtaining an F-value = 7.095 and p-value = 0.00951 **. The green line represents labor hours per month.

The need for editing or translation of scientific texts is widespread among Colombian doctoral students. Among the respondents, 93.9% have asked for favors to edit their English and 32.7% have asked for translation favors. Regarding the use of paid services, 59.2% have paid for editing their articles and 28.6% have paid for a translation. The *Premium* editing total cost and the standard translation cost represent almost a half of an average doctoral monthly salary in Colombia (Fig 2).

**Fig 2.**
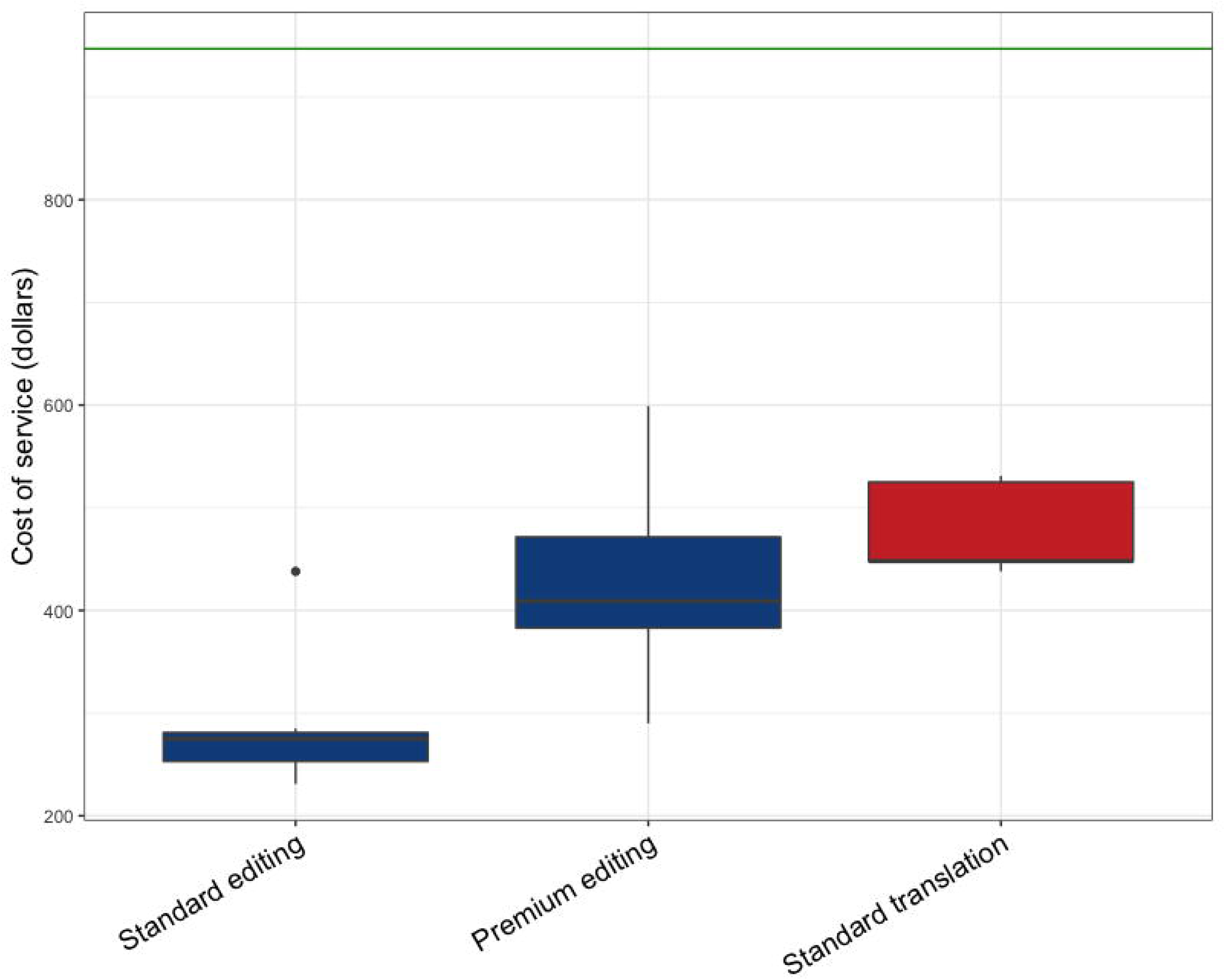
Cost of translation and editing services of scientific articles from six publishers. The Y axis is the price of the service in US dollars, the X axis represents the type of service, the standard or Premium service corresponds to the delivery days. The green line represents an average Ph.D salary in Colombia ($ 947).

Reading comprehension is also affected by the language of the text (Fig 3). However, only 18% of respondents prefer to read scientific articles in Spanish than in English. On the other hand, neither the interpretation of figures nor the understanding of scientific terminology is affected by the reading language.

**Fig 3.**
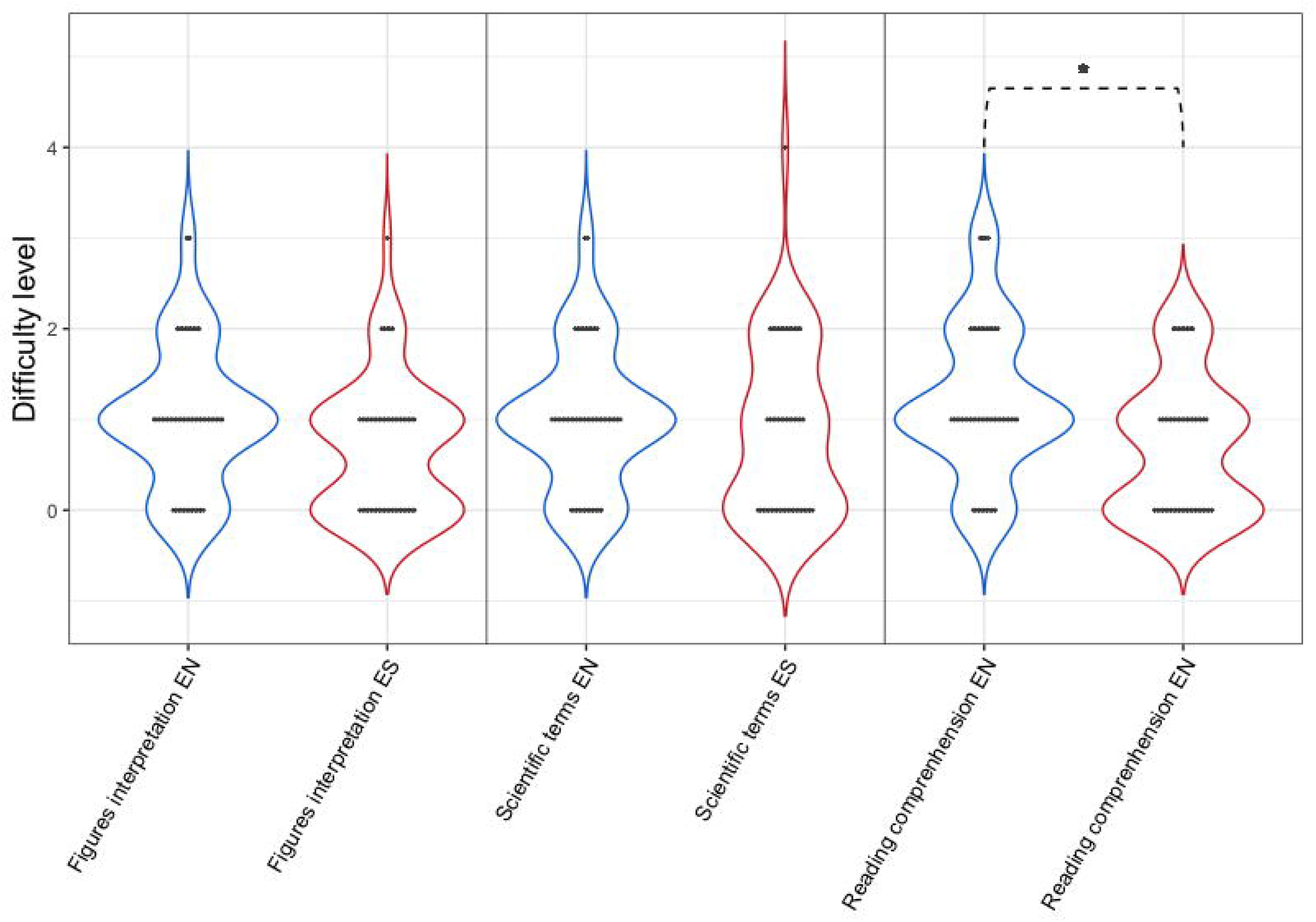
Reading comprehension, interpretation of figures, scientific terminology in English (EN) vs. Spanish (ES). A *Poisson* regression was used to analyze these discrete ordinal variants (Qualification from 1 to 5). A Chi-squared test was performed between languages for each category: interpretation of figures (Z-value = 0.756, Pr (Chi) = 0.09754), understanding of scientific terminology (z-value = 0.143, Pr (Chi) = 0.4619) and reading comprehension (z-value = 1.427, Pr (Chi) = 0.01209 *).

To analyze the difficulty of writing scientific articles in two languages, the most commonly found sections in an article were taken into account: introduction, methods, results, discussion and conclusions. In all cases, survey participants found the discussion was the most difficult section to write, while the methods were perceived as “easier” (Fig 4). Overall, all sections except methods are perceived as significantly “more difficult” to write in English than in the participant’s first language.

**Fig 4.**
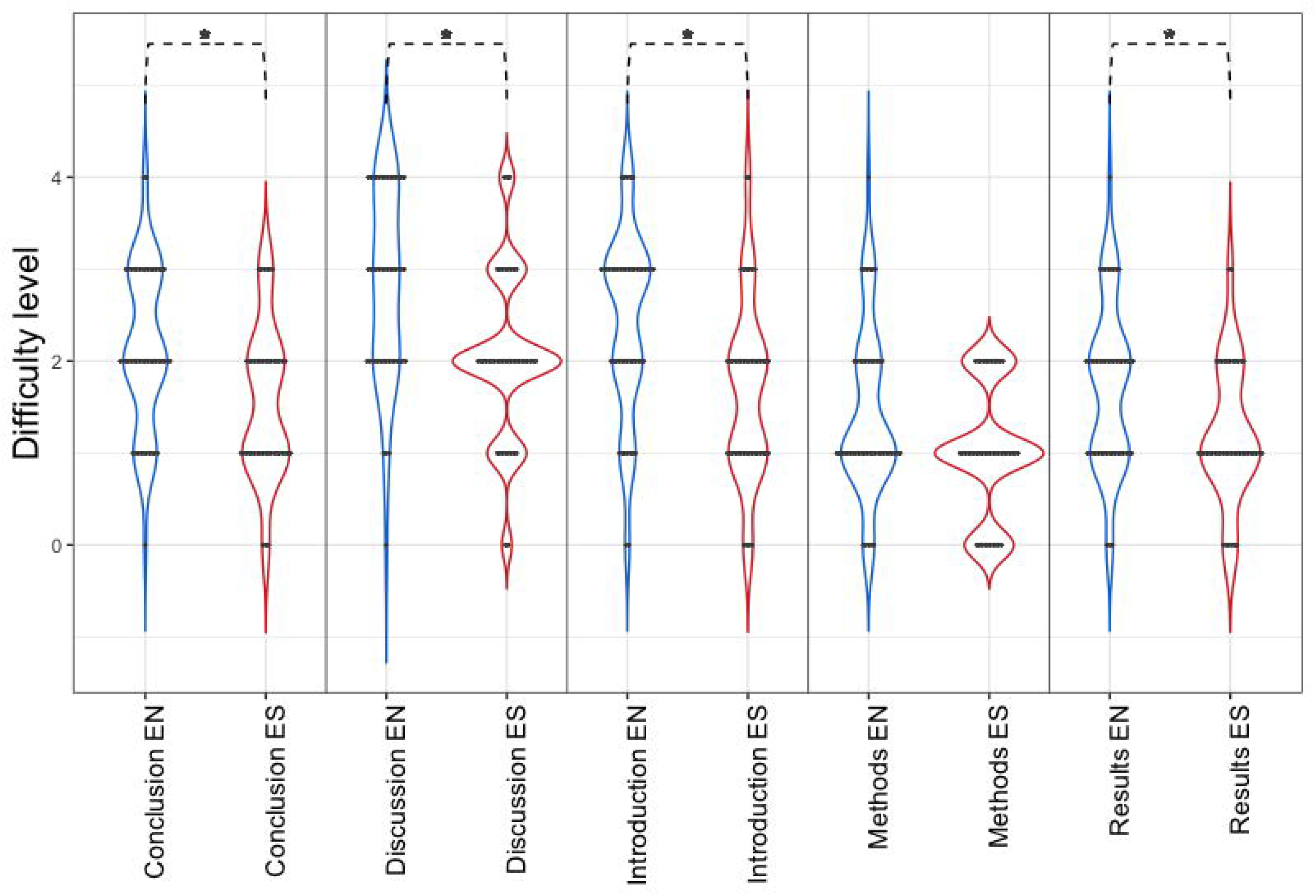
Difficulty of writing the different sections contained in a scientific article in English (EN) and Spanish (ES). A *Poisson* regression and Chi-square test was carried out: Introduction (z-value = 9.325, Pr (Chi) = 0.0158 *), methods (z-value = 3.046, Pr (Chi) = 0.07057), results (z-value = 4.899, Pr (Chi)= 0.04397 *), discussion (z-value = 11.732, Pr (Chi) = 0.02384 *), and conclusion (z-value = 7.688, Pr (Chi) = 0.03956 *).

With regard to the use of English in oral presentations at international events and conferences, 33% of respondents stated that they have stopped attending due to the mandatory use of English in oral presentations. Additionally, greater anxiety was perceived when presenting papers orally in English than in Spanish (Fig 5).

**Fig 5.**
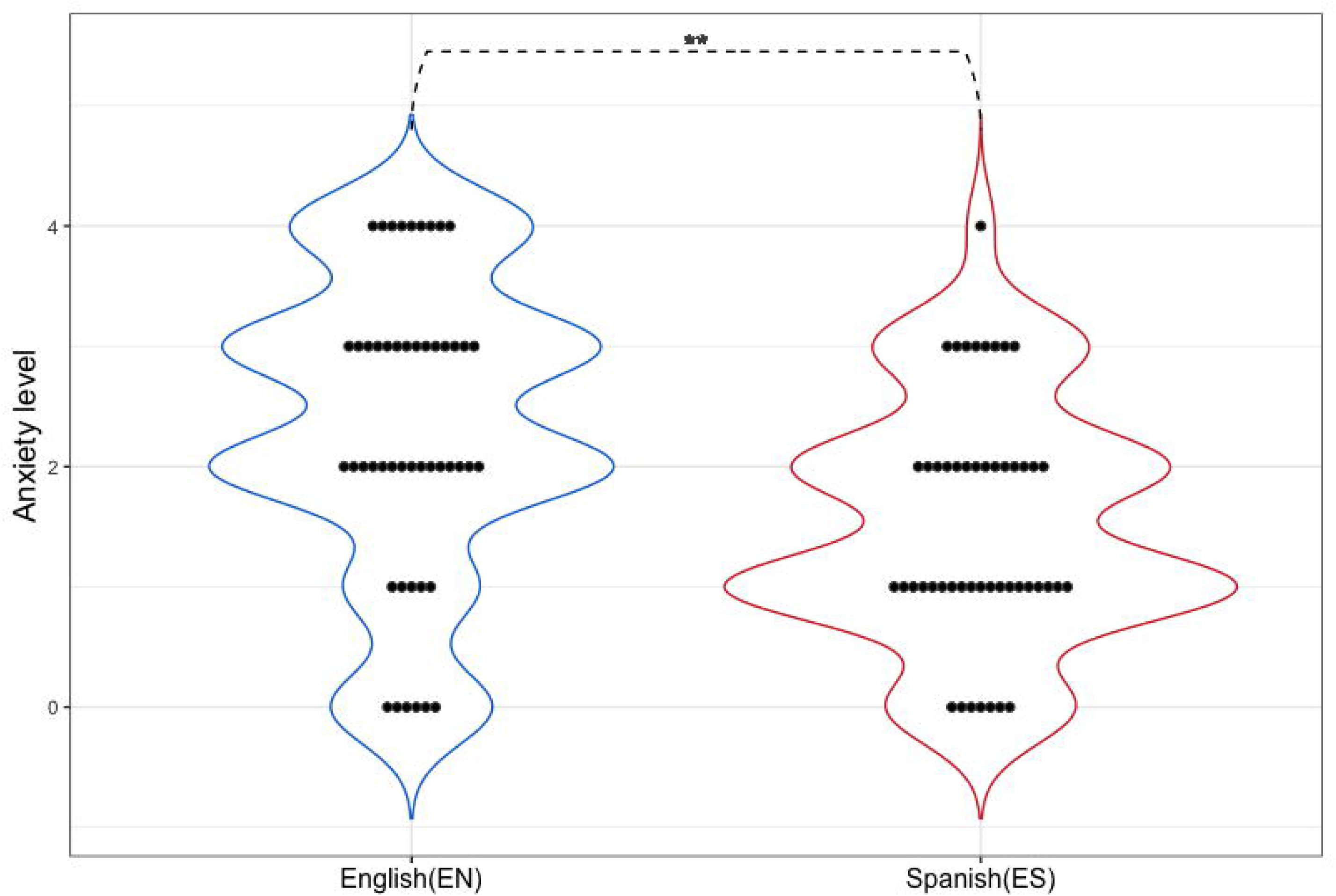
Anxiety level when making oral presentations in English (EN) vs. Spanish (ES). A *Poisson* regression was used to analyze discrete ordinal variants (Anxiety level from 1 to 5). A Chi-square test was carried out (z-value = 8,882, Pr (Chi) = 0.005419 **).

In order to determine whether or not the socioeconomic origin of doctoral students affects their proficiency in English and in turn increases the costs of publishing in English, an analysis of principal components was used reduce survey data related to socioeconomic background or English proficiency into single variables because both represent more than the 50% of the whole variance. For the following analyzes: 1) English proficiency is represented by PC1_English_proficiency, which explains 51% of the variance of the survey variables that are related to this subject (see methods), 2) the socioeconomic status is represented by PC1_Socioeconomic_status, which represents 62% of the variance of the variables of the survey that were related to this denomination (see methods). The socioeconomic status explains 15% of the English proficiency of researchers (Fig 6), which means that family and economic resources are partly translated into more proficient English.

**Fig 6.**
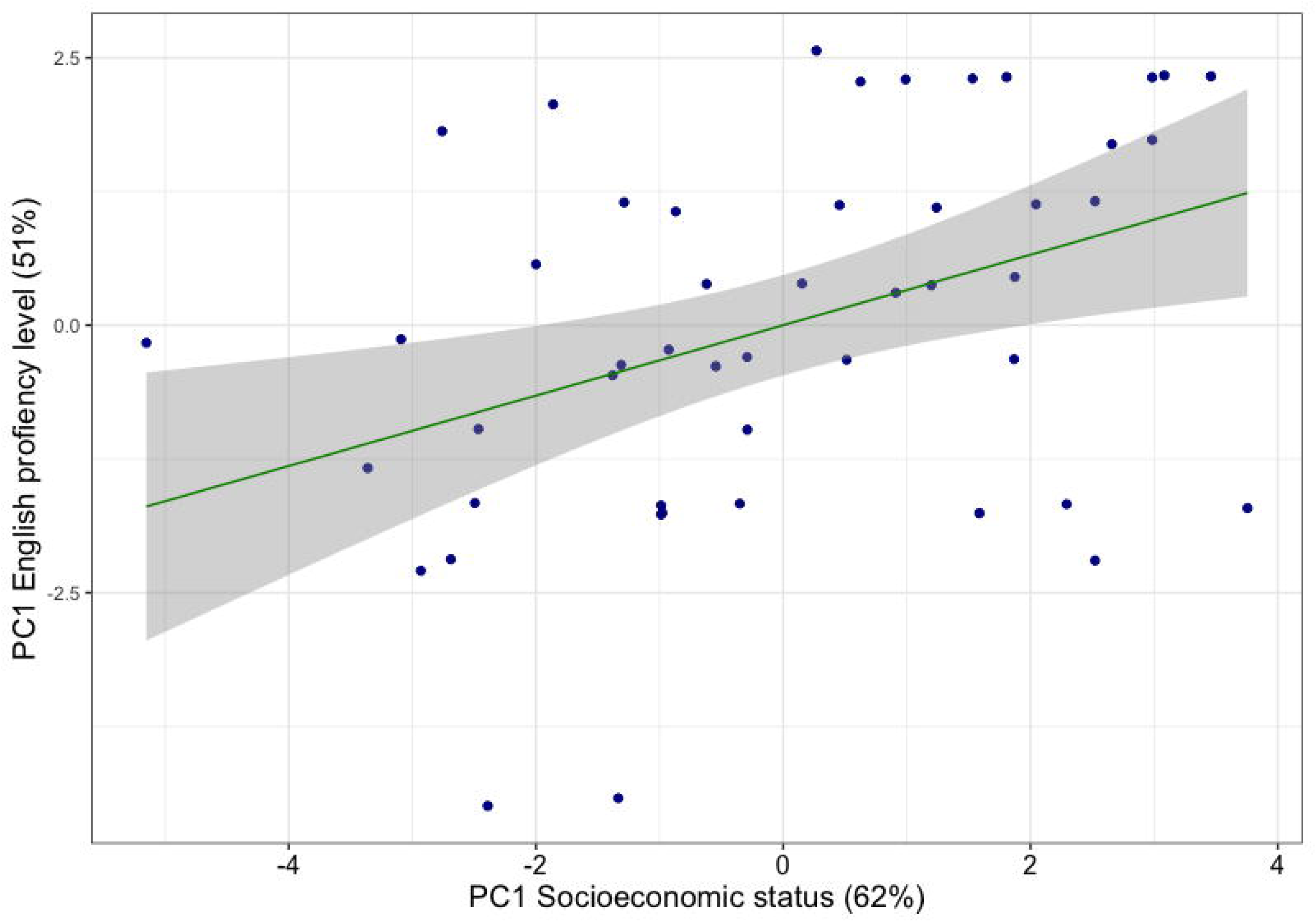
Relationship between socioeconomic status and English proficiency. Principle components representing socioeconomic status and English proficiency are significantly correlated (R2 = 0.1548, adjusted R2 = 0.1368, F = 8.605, p-value = 0.005168 **).

## Discussion

Many of the factors relating to publishing in English assessed in our study represent substantial costs in time, finances, productivity, and anxiety to Colombian researchers (Fig 7). Interestingly, the researchers appear to prefer to read and write articles in English and the scientific terminology do not represent an additional cost for Colombian researchers. In addition, a correlation between the socioeconomic status and English proficiency was found, suggesting an intersectional effect of language in science. These results can be extrapolated to understand costs of the English hegemony to all South American researchers, that in part contributes to a global gap between native English-speaking scientists (NES) and EFL scientists. This gap makes apparent the necessity of recognizing and protecting multilingualism in science. Although having common language is important for science communication, this effort should involve different actors in the research community and not only EFL researchers’ effort.

**Fig 7.**
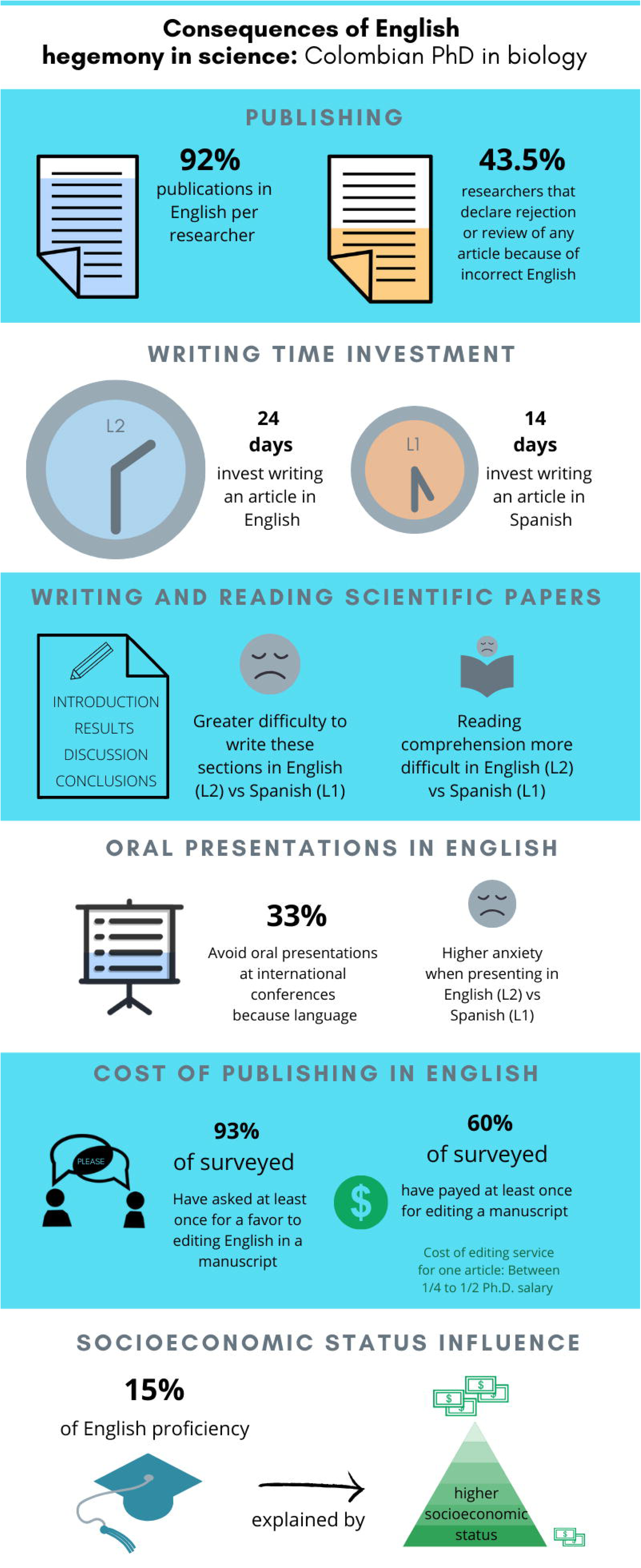
Summary of the results obtained organized by their impact on Colombia researchers. Second or foreign language (L2), first or native language (L1).

Our results show that several factors could lead to disadvantages of EFL researchers. The time investment in writing an article in English, for example, increases on average by 96.86 labor hours. This variable was not directly measured; it is based on the subjective perception of time of each person. However, as Guardiano and collaborators (2007) suggest, this extra cost affects the time spent on scientific tasks, decreasing the scientific productivity of researchers. Regarding the economic costs, between 50% and 30% of respondents have hired services to correct or translate scientific texts. To contextualize the cost of these services, a doctoral student should invest one-quarter to one-half doctoral monthly salary per article. It should be taken into account that scholarships and financing opportunities for doctoral students in the country are scarce (Fajardo de la Espriella, 2019), and not all of them have access to the forgivable loans provided by governmental institutions. More than 90% of researchers have asked for English-editing favors, but favors are a social cost. Social relations in the scientific world can have consequences on authorships of articles, collaborations with other laboratories, among others; this could reinforce dependence with research groups in NES countries (Flowerdew, 1999).

Around 80% of the respondents prefer to read and write scientific content directly in English. However, this result could be interpretable as “obligation” rather than as “preference” because of the monolingualism of scientific readings and the pressure to publish in international journals, and therefore in English (Guardiano et al, 2007). A scientist’s preference for reading and writing in English could also be due to the prevalence of English as the source for scientific words and phrases, as well as the scientist’s need to improve their own English in order to overcome these other barriers (Karimnia, 2013). The preference of writing directly in English and not translating may be related to the cost of the translation service that is three times the service for revision (Fig 2). Additionally, scientists are more likely to request a favor for English editing than for a translation (Guardiano et al., 2007). Strong feelings of insecurity or an “inferiority complex” generated by scientific writing in English is one of the most important segregation factors mentioned by EFL speaking researchers and increase the need of constant editing or correction (Flowerdew, 1999; Murasan & Pérez-Llantada, 2014; Huang, 2010). This difficulty or insecurity is augmented in the introduction and discussion sections of an article (Flowerdew, 2007; Burguess et al., 2014; Martín et al., 2014; Moreno & Rocha, 2012; Hanauer et al., 2019). However, the “materials and methods” section in an article and understanding scientific terminology are equally understood and used in both languages by the respondents, possibly because most words and expressions in modern science are coined in English (Ammon, 2001).

Coates (2002) shows that there is a greater probability of manuscript rejection by a journal if there are grammatical errors, and this study finds that such rejection has effects for EFL speaking researchers, since 43.5% of surveyed researchers reported suffering from rejection or revisions because of aspects related to grammar or style in English writing. However, without comparing this trend with native speakers it is not possible to conclude that this is worse for EFL researchers. Nevertheless, understanding reviewer comments is more difficult for a EFL speaking author, since these frequently contain expressions, euphemisms, or colloquialisms that are not easily interpreted by EFL speakers (Canagarajah, 2007; Fox & Canagarajah, 2008). For this reason, authors such as Huang (2010) call on reviewers to write comments that contribute and guide the use of English, and that do not discourage or criticize EFL authors for the lack of mastery of the language. On the other hand, “not every native English speaker is competent to solve peculiarities in the grammar and style of the “good” use of academic English”, therefore, all scientists have been pressured to use editing services (Tychinin & Webb 2003). In other words, it is questionable to judge or reject innovations or scientific research by linguistic factors or with the excuse of linguistic factors. If a particular research is important for the scientific community, the journal or other resources must assume the cost and effort of translation or editing services, shifting the costs from individual scientists to the publishers or the community.

It was expected that additional costs for Colombian researchers would be found, since similar findings have been reported from other EFL speaking countries in the world (McConnell, 1991; Flowerdew, 2007; Nour, 2005; Guardiano et al., 2007; Curry & Lillis, 2017; Hanauer et al., 2019). Despite the lack of specific studies on this subject across Latin America, with a few exceptions (Curry & Lillis, 2017; Hanauer et al., 2019), it is possible to assume that these results can be extrapolated to other countries bordering Colombia, given the similarity in proficiency and access to English, shared first language, low state investment in science and technology, and parallel political history with the US and Europe (Russell et al., 2008; British Council, 2015; Curry & Lillis, 2017). The results could even be extrapolated to other peripheral countries of the world, as Hanauer et al. (2019) found the similar disadvantages over doctoral students from two countries on different continents, Mexico and Taiwan. In addition, in this study we not only explore the impact that English proficiency has on doctoral students or post-doctoral researchers, but how those impacts are influenced by the researcher’s socioeconomic origin. A positive relationship (R^2^ = 0.14) was found between English proficiency and socioeconomic status, which is supported by previous studies (Fandiño-Parra, 2012), hence maintaining in science the patterns of social segregation at national and global levels.

This study finds that the system within science that denotes English as the lingua franca reinforces inequities between scientists from NES and EFL speaking countries, as well as socioeconomic inequities within countries that primarily speak a language other than English. Globalizing science, so far, has meant offering greater advantages to English speakers at the expense of other scientists’ prosperity in the world. Science at present, due to different pressures, opts for English as the only language acceptable for scientific communication, however, some researchers still value the protection of multilingualism in science (Bocanegra-Valle, 2014; Burguess et al., 2014). Defending multilingualism as an alternative in science would promote the reduction of international and social inequities, which would ultimately boost what Segatto (2019) has called “a radically plural world”. The homogenization of language in science with the excuse of “integration” is an expression of the elimination of diversity, and this can have consequences not only on the human diversity that makes science but on the diversity of scientific questions that arise (Alves & Pozzebonm, 2014).

The convenience of a common language in science must be recognized; however, it is essential that solutions to this problem involve scientists from a variety of backgrounds through a bilateral effort (EFL speaking scientists and NES speaking scientists) (Salager -Meyer, 2008; Muresan & Pérez-Llantada, 2014). Although research is a collective process, the proposed solutions so far have leaned on individual investment, which creates barriers to performing science that more greatly affect researchers of lower socioeconomic backgrounds. Universities, publishers, translation technology, conferences, among others, must also commit to generate ideas for change (Guardiano, 2007; Alves and Pozzebon, 2014). One potential approach would be to reduce the perceived value of publishing in journals with high impact factors (IFs), in order to reduce the pressure to publish in international journals (Murphy & Zhu, 2012; Rowley, 2017). Other alternatives include supporting journals that accept papers in several languages, promoting the inclusion of other languages in journals at the international level, incorporating revision or translation services in all fees paid to publish an article and providing these services to all scientists at no additional charge to them, establishing multilingual annual or periodic editions in renowned journals, among others (Guardiano, 2007). Proposals for universities and conferences include aids such as English tutoring for academic purposes (Universidad de los Andes Colombia, 2019), retaining in international conferences a space for presenting in local languages (Alves & Pozzebon, 2014), using methodologies such as simultaneous translation in conferences, and generating exchange spaces in other languages, among others. Finally, it would be helpful to strengthen public available technologies such as Google Translate that allow simultaneous written translation (Alves & Pozzebon, 2014). In the future, more alternatives will arise and it will be essential to analyze and monitor them to investigate their reception at the editorial and scientific level.

## Supporting information

S1 Complete article in Spanish.

S2 Summary of the results obtained organized by their impact on Colombia researchers in Spanish.

S3 Survey questions.

## Acknowledgements

Thanks to the researchers who completed the surveys or helped to share the survey. To Maria Carme Junyent Figueras for being the master thesis director that leads to this paper. To Pere Francesch Rom, Henry Arenas, Prof. Francesc Bernat, Prof. David Bueno and Prof. Avel·lí for editing and making suggestions on the original manuscript in Spanish. To the developers of Google Translate for creating a free powerful tool to translate in the first place the manuscript. To Rebecca Tarvin, Danny Jackson and Tyler Douglas and for editing and commenting on the manuscript in English. Finally, to the CRAI service of the University of Barcelona for providing the resources to publish this manuscript.

## Supplementary information

**S1 Complete article in Spanish**

**S2 Summary of the results obtained organized by their impact on Colombia researchers in Spanish.** Second or foreign language (L2), first or native language (L1).

**S3 Survey questions.** Questions in Spanish of the survey “Implications of language in scientific publications”.

