## Supplementary material for "Disadvantages of writing, reading, publishing and presenting scientific papers caused by the dominance of the English language in science: The case of Colombian PhD in biological sciences": S1 Complete article in Spanish.

### Desventajas de escribir, leer, publicar y presentar trabajos científicos a causa de la hegemonía del idioma inglés en la ciencia: el caso de los doctorados colombianos en ciencias biológicas

### Consecuencias de la hegemonía del inglés en la ciencia: el caso de los doctorados colombianos en ciencias biológicas

Valeria Ramírez-Castañeda[1,2,3]*

[1]Departamento de Filología, Universidad de Barcelona, Barcelona, España

[2]Departamento de Biología Integrativa, Universidad de California, Berkeley, Estados Unidos de América

[3]Museo de Zoología de Vertebrados, Universidad de California, Berkeley , Estados Unidos de América

* Autor para correspondencia

Correo electrónico: (VRC)

#### Resumen

El éxito de un científico depende tanto de su producción de artículos científicos como del factor de impacto de la revista donde son publicados. Debido a que las principales revistas científicas se publican en inglés, el éxito está relacionado con la publicación en el idioma anglosajón. Actualmente, el 98% de las publicaciones en ciencia están escritas en inglés, incluso entre investigadores de países con inglés como lengua extranjera. Colombia se encuentra entre los países con el nivel de inglés más bajo del mundo. Por lo tanto, comprender las desventajas que enfrentan los científicos en Colombia al producir artículos es crucial para reducir la desigualdad global en la ciencia. Este documento cuantifica las desventajas que resultan de la hegemonía del idioma en la publicación científica al examinar los costos adicionales que la comunicación en inglés crea en la producción de artículos científicos. Se identificó que más del 90% de los artículos científicos publicados por investigadores colombianos están en inglés, y que la publicación en un segundo idioma crea costos financieros adicionales para los estudiantes de doctorado colombianos. Además, genera problemas de comprensión de lectura y en la facilidad para escribir, y finalmente aumenta el tiempo de escritura de un artículo y la ansiedad a la hora de realizar presentaciones orales. El 43.5% de los estudiantes de doctorado reportó haber sufrido de rechazo o la revisión de sus artículos debido a la gramática del inglés, y el 33% eligió no asistir a conferencias internacionales debido al uso obligatorio del inglés en las presentaciones orales. Finalmente, entre los servicios de traducción/edición, el costo por artículo es entre un cuarto y la mitad de un salario mensual de un estudiante de doctorado en Colombia. De particular interés, identificamos una correlación positiva entre el dominio del inglés y el origen socioeconómico del investigador. En general, este estudio exhibe las consecuencias negativas de la hegemonía del inglés que preserva la brecha global en la ciencia. Aunque tener un idioma común es importante para la comunicación científica, generar alternativas desde el multilingüismo promovería la diversidad y al mismo tiempo conservaría un canal de comunicación. Tal esfuerzo debería provenir de diferentes actores y no debería recaer únicamente en los investigadores con inglés como idioma extranjero.

#### Introducción

Al mismo tiempo que los artículos científicos se convirtieron en la medida de la productividad científica, el inglés se impuso como el idioma de la ciencia, la cultura y la economía global (Johnson et al., 2018). Como consecuencia, hoy el 98% de las publicaciones en ciencias están escritas en inglés, especialmente en las áreas de ciencias naturales y básicas, estableciendo el inglés como la *lingua franca* de la ciencia (Gordin, 2015). Esto crea una desventaja para los científicos con inglés como idioma extranjero porque deben publicar textos complejos en un segundo idioma para avanzar en sus carreras (Parchomovsky, 1999). Esta desventaja da lugar a desigualdades globales, especialmente en países donde la mayoría de la población recibe una capacitación mínima en inglés y el bilingüismo con el inglés es muy bajo (Tardy, 2004). Por lo tanto, el dominio del inglés y el nivel socioeconómico influyen en el éxito científico, el acceso al conocimiento y la expatriación, entre otros.

Uno de los objetivos más importantes para la sociedad moderna es aumentar la producción científica de los paises del sur global (África, América Latina, Oriente Medio y de parte de Asia), dado que existe una fuerte correlación entre el dominio del inglés, el desarrollo económico y la innovación tecnológica en términos de número de artículos, número de investigaciones, y gastos de investigación y desarrollo (EF Education, 2018). Por lo tanto, la prevalencia del idioma inglés en las ciencias profundiza la desigualdad en la producción de conocimiento entre países con bajo dominio del inglés (Dei y Kempf, 2006), manteniendo la brecha en la producción científica entre los países del sur global o periférico y el países del norte global (incluidos los países del G8 y Australia), lo que reduce las contribuciones científicas individuales de los científicos con inglés como idioma extranjero (Whatmore, 2009, Murphy y Zhu, 2012).

Numerosos estudios han identificado el uso del inglés en la academia como una fuente de desigualdad y segregación en la ciencia (Flowerdew, 1999; Pabón-Escobar y da Costa, 2006; Muresan y Pérez-Llantada, 2014; Curry y Lilis, 2017, Hanauer et al. ., 2019). Estas inequidades afectan a la comunidad científica en múltiples niveles. En las comunidades locales de los países con inglés como idioma extranjero, el pensamiento científico se ve perjudicado, particularmente en la educación superior, ya que el aprendizaje depende de actitudes culturales derivadas del idioma nativo y la educación en ciencia se vuelve ajena a sus propias experiencias (Gulbrandsen et al., 2002; Lee &

Fradd, 2007; de Vasconcelos, 2006). La diversidad en el idioma promueve la diversidad en el pensamiento, afectando el proceso creativo y la imaginación; por lo tanto, el mantenimiento del multilingüismo en la ciencia podría tener un impacto en el conocimiento científico en sí mismo (Lee y Fradd, 2007).

Las revistas locales son un refugio para la comunicación de la investigación científica en otros idiomas además del inglés, sin embargo, a menudo se perciben como de baja calidad, ya que el trabajo de investigación más importante a menudo se reserva para revistas internacionales. Por lo tanto, los lectores con barreras idiomáticas solo tienen acceso a este tipo de conocimiento e incluso desconocen la investigación más importante que se está llevando a cabo en su propia región, lo que ha resultado en un vacío de información importante para la toma de decisiones políticas, las políticas ambientales regionales y las estrategias de conservación dentro de los territorios, derivando en un sesgo para la toma de decisiones a nivel mundial (Salager-Meyes, 2006; Alves y Pozzebonm, 2014; Amano et al., 2016). Además, a pesar de la importancia del conocimiento local, el éxito profesional de un científico se correlaciona en mayor medida con su "internacionalización". Esta presión constante podría estar influyendo en la migración académica, conocida como "expatriación". El aprendizaje del inglés es uno de los factores de presión migratoria, ya que es más difícil lograr un dominio superior del inglés para los científicos que permanecen en los países con inglés como idioma extranjero (Benfield y Howard, 2000; Ferguson, 1999; de Vasconcelos, 2006).

En los países de la periferia existe una fuerte relación entre el dominio del inglés y el origen socioeconómico, por lo que es importante comprender los costos de publicación asociados con el origen socioeconómico del estudiante de doctorado. Entre el sur global, Colombia es el segundo territorio más desigual en Latinoamérica: en 2018 invirtió sólo el 0.24% de su PIB (la inversión de Suecia fue el 2.74% de su PIB) en ciencia, tecnología e innovación (Portafolio, 2019), y tiene uno de los niveles más bajos de dominio del inglés entre los rankings mundiales (EF Education, 2018). Además, solo el 0.015% de la población tiene o está haciendo un doctorado en ciencias básicas o naturales (UNESCO, 2016). Este estudio tiene como objetivo determinar si los estudiantes de doctorado colombianos en ciencias naturales enfrentan desventajas al publicar artículos científicos en inglés, en comparación con las publicaciones en su primer idioma, y cuantificar el trabajo adicional que estos científicos realizan en la escritura, lectura y presentación de su trabajo en inglés. Además, este estudio examina el impacto del origen socioeconómico en el dominio del inglés y los costos que genera al publicar.

#### Materiales y métodos

Con el fin de determinar los costos de publicar en inglés experimentados por estudiantes de doctorado en ciencias biológicas, 49 estudiantes de doctorado colombianos participaron en la encuesta "Implicaciones del lenguaje en publicaciones científicas" que contiene 44 preguntas en idioma español (preguntas de la encuesta S3) . Además, se buscaron los precios ofrecidos por prestigiosas editoriales científicas para la traducción (español al inglés) y la edición de textos científicos para medir el impacto económico en relación con un salario promedio de doctorando Colombia.

#### Construcción de la encuesta

La encuesta principal de este trabajo, titulada “Implicaciones del lenguaje en las publicaciones científicas”, contiene 44 preguntas divididas en tres secciones: datos básicos, redacción de artículos en inglés y aprendizaje del inglés (preguntas de la encuesta S3). Las respuestas obtenidas se agruparon para el análisis estadístico (Tabla 1).

**Tabla 1.** Resumen de las preguntas de la encuesta divididas por grupos utilizados para el análisis estadístico correspondiente.

| Grupo | Variable | Tipo de variable | Referencia |
|---|---|---|---|
| **Datos socioeconómicos** | Asistir a educación pública o privada en el bachillerato y la universidad | Binario (público / privado) | Cataño, 2017 |
| | Ocupación del padre y la madre para definir el origen socioeconómico | Categórica | Requena, 2012 |
| | Estratificación socioeconómica en Colombia de él - o ella misma, padre y madre | Ordinal (1 a 6) | El Congreso de Colombia, 1994 |
| **Dominio del inglés** | Primer idioma (s) | Categórico | |
| | Idioma del país en el que viven. | Categórico | |
| | Competencia en escritura, expresión oral, lectura y comprensión oral en inglés | Ordinario (malo, regular, bueno, excelente) | |
| | Edad de inicio del aprendizaje del inglés Cuantitativo | Discreto | |
| | Años estudiando inglés | Cuantitativo discreto | |
| | Experiencia satisfactoria en la institución de aprendizaje de inglés | Ordinal (1 a 5) | |
| | Años vividos en el país de habla inglesa | Discreto cuantitativo | |
| | Porcentaje de inglés que habla diariamente | Porcentaje cuantitativo | |
| **Publicaciones** | Número de publicaciones científicas revisadas por pares académicos | Cuantitativo discreto | |
| | Porcentaje de esas publicaciones en inglés o en un idioma que no sea inglés | Porcentaje cuantitativo | |

| | | |
|---|---|---|
| **Costos de escribir en inglés** | Pago o favores para editar o traducir publicaciones científicas | Binario (sí / no) |
| | Porcentaje de artículos: pago o favores por editar o traducir publicaciones científicas | Porcentaje cuantitativo |
| | Número de rechazos o revisiones en revistas científicas asociadas a la escritura en inglés | Discreto cuantitativo |
| | Tiempo dedicado a escribir un artículo científico en inglés o español (horas laborales) | Cuantitativo continuo |
| | Preferencia entre escribir directamente en inglés o traducir al inglés | binario (inglés / traducir) |
| **Dificultad de escritura en inglés o español** | Dificultad para escribir la sección de introducción , métodos, resultados, discusión o conclusiones de un artículo científico | Ordinal (1 a 5) |
| **Dificultad de lectura en inglés o español** | Dificultad para entender textos generales / Comprender terminología científica / Interpretar figuras y tablas | Ordinal (1 a 5) |
| **Participación en conferencias** | Presentación oral en conferencias internacionales | Binario (sí / no) |
| | Nivel de ansiedad en presentaciones orales en inglés o español | ordinal (1 a 5) |

#### Análisis estadístico

Los análisis estadísticos se realizaron en R v.3.6.1 (R Development Core Team, 2011) y los datos se graficaron con el paquete ggplot (Valero-Mora, 2015). Para comparar la lectura y escritura entre inglés y español, el tiempo invertido en escribir y el nivel de ansiedad en la participación en conferencias, se realizó un ANOVA (*aov* en el paquete 'stats' v3.5.3). El margen de error se calculó con un 95% de confianza. Se realizó un Análisis de Componentes Principales (PCA) utilizando las variables contenidas en los grupos "inglés" y "Datos socioeconómicos" para reducir la redundancia en las variables (*PCA* en el paquete Dominio delFactorMiner v2.2). Se revisó la proporción de varianza explicada por cada componente principal, y solo se retuvo el primer componente principal para cada conjunto de datos, ya que describió el 51% y el 62% correspondientemente de la variación total. Posteriormente, se ejecutó una regresión lineal con la intención de comparar estas dos variables, el dominio del inglés PC1 versus el nivel socioeconómico PC1 (usando *lm* en el paquete 'stats' v3.5.3).

#### Costos de los servicios de edición y traducción

Para visualizar los precios de los servicios de edición y traducción en inglés de textos científicos, se buscó información en cinco de las editoriales científicas más importantes (SAGE language services, 2018; Cambridge University Press Author Services, 2018; Elsevier author services). , 2018; SpringerNature author services, 2018 y Wiley edition services, 2018). La información y los costos de estos servicios son públicos y se pueden obtener a través de las páginas web de los editores. Todos los datos se tomaron con respecto a los precios de un texto de 3000 palabras, ya que esa es la longitud promedio de un artículo científico; Las búsquedas se realizaron en octubre de 2018.

Estas editoriales ofrecen dos tipos de servicios de edición, un servicio de tres días (Premium) y un servicio de una semana (estándar); ambos precios fueron utilizados para el análisis. Solo se utilizaron los precios de las traducciones español - inglés. Finalmente, estos precios se compararon con un salario doctoral promedio en Colombia (Colciencias, 2019), 947 dólares estadounidenses o 3 millones de pesos colombianos (1 dólar estadounidense = 3.166 pesos colombianos, precio de cambio al 31 de enero de 2019).

#### Resultados

Se obtuvieron un total de 49 respuestas de estudiantes de doctorado o doctorandos colombianos en ciencias biológicas cuyo primer idioma es el español. De los colombianos encuestados, el 92% (sd = 0.272) de sus artículos científicos publicados están en inglés y solo el 4% (sd = 0.2) de sus publicaciones estaban en español o portugués. Además, el 43.5% de los estudiantes de doctorado declararon al menos un rechazo o revisión de sus artículos debido a la gramática del inglés.

Con respecto a la inversión de tiempo, hubo un aumento significativo en el tiempo invertido escribiendo un artículo científico en inglés en comparación con el español para los participantes de la encuesta (Fig. 1). El proceso de escritura en español toma en promedio 114.57 (sd = 87.77) horas laborales, mientras que en inglés, 211.4 (sd = 182.6) horas laborales. En promedio, estos científicos pasan 96.86 horas laborales más escribiendo en inglés. Sin embargo, el 81.2% de los estudiantes de doctorado declararon que prefieren escribir directamente en inglés en comparación con escribir en español y luego traducir al inglés.

**Fig. 1.** Tiempo invertido escribiendo un artículo científico en inglés (EN) como idioma extranjero y en español (ES) como primer idioma. Se realizó un análisis ANOVA para comparar las variables obteniendo un valor $F = 7.095$ y un valor $p = 0.00951$ **. La línea verde representa las horas laborales por mes.

La necesidad de editar o traducir textos científicos está muy extendida entre los estudiantes de doctorado colombianos. Entre los encuestados, el 93.9% ha pedido favores para editar su inglés y el 32.7% ha pedido favores de traducción. Con respecto al uso de servicios pagos, el 59.2% ha pagado por editar sus artículos y el 28.6% ha pagado por una traducción.

El costo total de edición tipo *premium* y el costo de traducción estándar representan casi la mitad de un salario mensual promedio de doctorado en Colombia (Fig. 2).

**Fig. 2.** Costo de los servicios de traducción y edición de artículos científicos de seis editoriales científicas. El eje Y es el precio del servicio en dólares estadounidenses, el eje X representa el tipo de servicio, el servicio estándar o *Premium* corresponde a los días de entrega. La línea verde representa un salario promedio de doctorado en Colombia ($ 947).

La comprensión de lectura también se ve afectada por el lenguaje del texto (Fig. 3). Sin embargo, solo el 18% de los encuestados prefiere leer artículos científicos en español que en inglés. Por otro lado, ni la interpretación de las figuras ni la comprensión de la terminología científica se ven afectadas por el lenguaje de lectura.

**Fig. 3.** Comprensión de lectura, interpretación de figuras, terminología científica en inglés (EN) vs. español (ES). Se utilizó una regresión de *Poisson* para analizar estas variables ordinales discretas (Calificación del 1 al 5). Se realizó una prueba de Chi cuadrado entre idiomas para cada categoría: interpretación de figuras (valor Z = 0.756, Pr (Chi) = 0.09754), comprensión de la terminología científica (valor z = 0.143, Pr (Chi) = 0.4619) y comprensión de lectura (valor z = 1.427, Pr (Chi) = 0.01209 *).

Para analizar la dificultad de escribir artículos científicos en dos idiomas, se tuvieron en cuenta las secciones más comúnmente encontradas en un artículo: introducción, métodos, resultados, discusión y conclusiones. En todos los casos, los participantes de la encuesta encontraron que la discusión era la sección más difícil de escribir, mientras que los métodos se percibían como "más fáciles" (Fig. 4). En general, todas las secciones, excepto los métodos, se perciben como significativamente "más difíciles" de escribir en inglés que en el primer idioma del participante.

**Fig. 4.** Dificultad para escribir las diferentes secciones contenidas en un artículo científico en inglés (EN) y español (ES). Se utilizó una regresión de *Poisson* y un análisis de Chi-cuadrado: Introducción (valor z = 9.325, Pr (Chi) = 0.0158 *), métodos (valor z = 3.046, Pr (Chi) = 0.07057), resultados (valor z = 4.899, Pr (Chi) = 0.04397 *), discusión (valor z = 11.732, Pr (Chi) = 0.02384 *) y conclusión (valor z = 7.688, Pr (Chi) = 0.03956 *).

Con respecto al uso del inglés en presentaciones orales en eventos y conferencias internacionales, el 33% de los encuestados declararon que habían dejado de asistir debido al uso obligatorio del inglés en las presentaciones orales. Además, se percibió una mayor ansiedad al presentar trabajos en inglés que en español (Fig. 5).

**Fig. 5.** Nivel de ansiedad al hacer presentaciones orales en inglés (EN) vs. español (ES). Se utilizó una regresión de *Poisson* para analizar variables ordinales discretas (nivel de ansiedad de 1 a 5). Se realizó una prueba de Chi-cuadrado (valor z = 8,882, Pr (Chi) = 0,005419 **).

Para determinar si el origen socioeconómico de los estudiantes de doctorado afecta o no su dominio del inglés y, a su vez, aumenta los costos de publicación en inglés, se utilizó un Análisis de los Componentes Principales (PCA) para reducir las variables de la encuesta relacionados con los antecedentes socioeconómicos o el dominio del inglés en variables únicas que representan más del 50% de toda la varianza. Para los siguientes análisis: 1) El dominio del inglés está representado por PC1_English_proficiency, que explica el 51% de la varianza de las variables de la encuesta relacionadas con este tema (ver métodos), 2) el estado socioeconómico está representado por PC1_Socioeconomic_status, que representa el 62% de la varianza de las variables de la encuesta que están relacionadas con esta denominación (ver métodos). El origen socioeconómico explica el 15% del dominio del inglés de los investigadores (Fig. 6), lo que significa que los recursos familiares y económicos se traducen en parte en un mayor nivel de inglés.

**Fig 6.** Relación entre el estatus socioeconómico y el dominio del inglés. Los componentes principales que representan el estatus socioeconómico y el dominio del inglés están significativamente correlacionados ($R2 = 0.1548$, R2 ajustado = 0.1368, $F = 8.605$, valor $p = 0.005168$ **).

#### Discusión

Muchos de los factores relacionados con la publicación en inglés evaluados en nuestro estudio representan costos sustanciales en tiempo, finanzas, productividad y ansiedad para los investigadores colombianos (Figura 7). Curiosamente, los investigadores parecen preferir leer y escribir artículos en inglés y la terminología científica no representa un costo adicional para los investigadores colombianos. Además, se encontró una correlación entre el nivel socioeconómico y el dominio del inglés, lo que sugiere un efecto interseccional del lenguaje en la ciencia. Estos resultados se pueden extrapolar para comprender los costos de la hegemonía de inglés para todos los investigadores sudamericanos, lo que en parte contribuye a una brecha global entre los científicos nativos de habla inglesa y los científicos con inglés como lengua extranjera. Esta brecha pone de manifiesto la necesidad de reconocer y proteger el multilingüismo en la ciencia. Aunque tener un lenguaje común es importante para la comunicación científica, este esfuerzo debe involucrar a diferentes actores en la comunidad de investigación y no solo el esfuerzo de los investigadores con inglés como lengua extranjera.

**Fig 7. Resumen de los resultados obtenidos organizados por su impacto en los investigadores colombianos.** Segundo idioma o idioma extranjero (L2), primer idioma o lengua materna (L1).

Nuestros resultados muestran que varios factores podrían conducir a desventajas de los investigadores con inglés como lengua extranjera. El tiempo invertido en escribir un artículo en inglés, por ejemplo, aumenta en promedio en 96.86 horas laborales. Esta variable no se midió directamente, se basa en la percepción subjetiva del tiempo de cada persona. Sin embargo, como sugieren Guardiano y colaboradores (2007), este costo adicional afecta el tiempo dedicado a tareas científicas, disminuyendo la productividad científica de los investigadores. En cuanto a los costos económicos, entre el 50% y el 30% de los encuestados han contratado servicios para corregir o traducir textos científicos. Para contextualizar el costo de estos servicios, un estudiante de doctorado debe invertir entre un cuarto y medio salario mensual completo de doctorado por artículo. Debe tenerse en cuenta que las becas y las oportunidades de financiamiento para estudiantes de doctorado en el país son escasas (Fajardo de la Espriella, 2019), y no todas estos tienen acceso a los préstamos condonables otorgados por instituciones gubernamentales. Más del 90% de los investigadores han pedido favores de edición en inglés, pero los favores son un costo social. Las relaciones sociales en el mundo científico pueden tener consecuencias sobre la autoría de artículos, colaboraciones con otros laboratorios, entre otros; esto podría reforzar la dependencia con los grupos de investigación en los países con inglés como idioma nativo (Flowerdew, 1999).

Alrededor del 80% de los encuestados prefieren leer y escribir contenido científico directamente en inglés. Sin embargo, este resultado podría interpretarse como "obligación" más que como "preferencia" debido al monolingüismo predominante de las lecturas científicas y la presión de publicar en revistas internacionales y, por lo tanto, en inglés (Guardiano et al, 2007). La preferencia de un científico por leer y escribir en inglés también podría deberse a la prevalencia del inglés como fuente de terminología científica, así como a la necesidad del científico de mejorar su propio inglés para superar estas otras barreras (Karimnia, 2013) . La preferencia de escribir directamente en inglés y no traducir puede estar relacionada con el costo del servicio de traducción que es tres veces el servicio de revisión (Figura 2). Además, es más probable que los científicos soliciten un favor para la edición en inglés que para una traducción (Guardiano et al., 2007). Los fuertes sentimientos de inseguridad o "complejo de inferioridad" generado por la escritura científica en inglés es uno de los factores de segregación más importantes mencionados por los investigadores que hablan inglés como lengua extranjera y aumentan la necesidad de una edición o corrección constante (Flowerdew, 1999; Murasan & Pérez-Llantada, 2014 ; Huang, 2010). Esta dificultad o inseguridad se incrementa en las secciones de introducción y discusión de un artículo (Flowerdew, 2007; Burguess et al., 2014; Martín et al., 2014; Moreno & Rocha, 2012; Hanauer et al., 2019). Sin embargo, la sección de "materiales y métodos" en un artículo y la comprensión de la terminología científica son igualmente entendidos y utilizados en ambos idiomas por los encuestados, posiblemente porque la mayoría de las palabras y expresiones en la ciencia moderna están acuñadas en inglés (Ammon, 2001).

Coates (2002) mencionó que existe una mayor probabilidad de rechazo de manuscritos por parte de una revista si hay errores gramaticales, y este estudio encuentra que dicho rechazo tiene efectos para los investigadores que hablan inglés como idioma extranjero, ya que el 43.5% de los investigadores encuestados reportaron sufrir rechazo o revisiones por aspectos relacionados con la gramática o el estilo en la escritura en inglés. Sin embargo, sin comparar esta tendencia con investigadores con inglés como idioma nativo, no es posible concluir que esto es peor para los investigadores con inglés como segunda lengua. Sin embargo, comprender los comentarios de los revisores es más difícil para un autor con inglés no nativo, ya que con frecuencia contienen expresiones, eufemismos o coloquialismos que no pueden interpretar fácilmente (Canagarajah, 2007; Fox y Canagarajah, 2008). Por esta razón, autores como Huang (2010) solicitan a los revisores que escriban comentarios que contribuyan y guíen el uso del inglés, y que no desalienten ni critiquen a los autores por la falta de dominio del idioma. Por otro lado, "no todos los hablantes nativos de inglés son competentes para resolver peculiaridades en la gramática y el estilo del buen uso del inglés académico", por lo tanto, todos los científicos están presionados a usar los servicios de edición (Tychinin y Webb 2003). En otras palabras, es cuestionable juzgar o rechazar las innovaciones o la investigación científica por factores lingüísticos o con la excusa de factores lingüísticos. Si una investigación en particular es importante para la comunidad científica, la revista u otros recursos deben asumir el costo y el esfuerzo de los servicios de traducción o edición, transfiriendo los costos de los científicos individuales a los editores o la comunidad.

Era de esperarse que se iban a encontrar costos adicionales para los investigadores colombianos, ya que se han reportado hallazgos similares de otros países con inglés como idioma extranjero en el mundo (McConnell, 1991; Flowerdew, 2007; Nour, 2005; Guardiano et al., 2007; Curry y Lillis , 2017; Hanauer et al., 2019). A pesar de la falta de estudios específicos sobre este tema en América Latina, con algunas excepciones (Curry y Lillis, 2017; Hanauer et al., 2019), es posible suponer que estos resultados se pueden extrapolar a otros países limítrofes con Colombia, dada la similitud en el dominio y el acceso al inglés, el primer idioma compartido, la baja inversión estatal en ciencia y tecnología y la historia política paralela con los Estados Unidos y Europa (Russell et al., 2008; British Council, 2015; Curry & Lillis, 2017). Los resultados podrían incluso extrapolarse a otros países periféricos del mundo, Hanauer et al. (2019) encontraron desventajas similares sobre los estudiantes de doctorado de dos países en diferentes continentes, México y Taiwán. Además, en este estudio no solo exploramos el impacto que el dominio del inglés tiene en los estudiantes de doctorado o doctorandos, sino también cómo esos impactos están influenciados por el origen socioeconómico del investigador. Existe una relación positiva ($R^2$encontró= 0.14) entre el dominio del inglés y el nivel socioeconómico, lo cual es respaldado por estudios previos (Fandiño-Parra, 2012), por lo tanto, en la ciencia se mantienen los patrones de segregación social a nivel nacional y global.

Este estudio encuentra que el sistema en la ciencia que mantiene al inglés como la *lingua franca* refuerza las desigualdades entre los científicos provenientes de países con alto nivel de inglés en comparación con los provenientes de países con nivel bajo, así como se amplían las desigualdades socioeconómicas dentro de los países que hablan principalmente un idioma que no es inglés. Globalizar la ciencia, hasta ahora, ha significado ofrecer mayores ventajas a los angloparlantes a expensas de la prosperidad de otros científicos en el mundo. La ciencia en la actualidad, debido a las diferentes presiones, opta por el inglés como el único idioma aceptable para la comunicación científica, sin embargo, algunos investigadores aún valoran la protección del multilingüismo en la ciencia (Bocanegra-Valle, 2014; Burguess et al., 2014). Defender el multilingüismo como alternativa en la ciencia promovería la reducción de las desigualdades internacionales y sociales, lo que en última instancia impulsaría lo que Segatto (2019) ha llamado "un mundo radicalmente plural". La homogeneización del lenguaje en la ciencia con la excusa de la "integración" es una expresión de la eliminación de la diversidad, y esto puede tener consecuencias no solo en la diversidad humana que hace ciencia sino en la diversidad de preguntas científicas que surgen (Alves y Pozzebonm, 2014).

Se debe reconocer la conveniencia de un lenguaje común en la ciencia; sin embargo, es esencial que las soluciones a este problema involucren a científicos de diversos orígenes a través de un esfuerzo bilateral (científicos que hablan inglés como lengua nativa y científicos que no) (Salager -Meyer, 2008; Muresan & Pérez-Llantada, 2014). Aunque la investigación es un proceso colectivo, las soluciones propuestas hasta ahora se han recostado en el esfuerzo individual, lo que crea barreras para la realización de la ciencia que afectan en mayor medida a los investigadores de entornos socioeconómicos más bajos. Las universidades, editoriales, tecnologías de traducción, conferencias, entre otros, también deben comprometerse a generar ideas para el cambio (Guardiano, 2007;

Alves y Pozzebon, 2014). Un enfoque potencial sería reducir el valor percibido de publicar en revistas con factores de alto impacto (IF), a fin de reducir la presión para publicar en revistas internacionales (Murphy y Zhu, 2012; Rowley, 2017). Otras alternativas incluyen el apoyo a revistas que aceptan trabajos en varios idiomas, promoviendo la inclusión de otros idiomas en revistas a nivel internacional, incorporando servicios de revisión o traducción gratuitos para publicar un artículo, estableciendo ediciones multilingües anuales o periódicas en revistas de renombre, entre otras (Guardiano, 2007). Las propuestas para universidades y conferencias incluyen ayudas como la tutoría de inglés con fines académicos (Universidad de los Andes Colombia, 2019), mantener en congresos internacionales un espacio para presentar en los idiomas locales (Alves y Pozzebon, 2014), utilizando metodologías como la traducción simultánea en conferencias, y generando espacios de intercambio en otros idiomas, entre otros. Finalmente, sería útil fortalecer las tecnologías disponibles públicamente, como Google Translate, que permiten la traducción escrita simultánea (Alves y Pozzebon, 2014). En el futuro, surgirán más alternativas y será esencial analizarlas y monitorearlas para investigar su recepción a nivel editorial y científico.

#### Agradecimientos

Gracias a los investigadores que completaron las encuestas o ayudaron a compartir la encuesta. A Maria Carme Junyent Figueras por ser la directora de tesis de maestría que conllevo a este proyecto. A Pere Francesch Rom, Henry Arenas, Prof. Francesc Bernat, Prof. David Bueno y Prof. Avel·lí por editar y hacer sugerencias sobre el manuscrito original en español. A los desarrolladores de Google Translate por crear una poderosa herramienta gratuita para traducir en primer lugar el manuscrito. A Rebecca Tarvin,

Danny Jackson y Tyler Douglas y por editar y comentar el manuscrito en inglés. Finalmente, al servicio CRAI de la Universidad de Barcelona por proporcionar los recursos para publicar este manuscrito.

#### Referencias

1. Alves MA, Pozzebon M. ¿Cómo resistir la dominación lingüística y promover la diversidad del conocimiento? Rev Adm Empres. 2014; 53: 629–633. doi: 10.1590 / s0034-759020130610

2. Amano T, González-Varo JP, Sutherland WJ. Los idiomas siguen siendo una barrera importante para la ciencia global. PLoS Biol. 2016; 14: e2000933. doi: 10.1371 / journal.pbio.2000933

3. Ammon U. El dominio del inglés como idioma de la ciencia: efectos en otros idiomas y comunidades lingüísticas. Mouton de Gruyter; 2001.

4. Benfield J, Howard K. El lenguaje de la ciencia. Eur J Cardio-Torácica Surg. 2000; 18: 642–648. doi:10.1016/S1010-7940(00)00595-9

5. Bocanegra-Valle A. “English is my default academic language”: Voices from LSP scholars publishing in a multilingual journal. J English Acad Purp. 2014;13: 65–77. doi:10.1016/j.jeap.2013.10.010

6. British Council. English in Colombia: An examination of policy, perceptions and influencing factors. 2015.

12. Curry MJ, Lillis TM. Global academic publishing : policies, perspectives and pedagogies. Bristol: Blue Ridge Summit; 2017.

13. de Vasconcelos Hage SR, Cendes F, Montenegro MA, Abramides DV, Guimarães CA, Guerreiro MM. Specific language impairment: linguistic and neurobiological aspects. Arq Neuropsiquiatr. 2006;64: 173–80. doi:/S0004-282X2006000200001

14. Dei GJS (George JS, Kempf A. Anti-colonialism and education : the politics of resistance. Sense Publishers; 2006.

15. EF Education. EF EPI 2018 - EF English Proficiency Index - Europe. In: EF Education First [Internet]. 2018 [cited 9 Jan 2019]. Available: https://www.ef.com.es/epi/

16. El Congreso de Colombia. Ley 142 de 1994. Ley 142 de 1994. D Of. 1994;

29. Husson F, Josse J, Le S, Mazet J. FactoMineR: Multivariate Exploratory Data Analysis and Data Mining with R. 2013.

36. McConnell GD. A macro-sociolinguistic analysis of language vitality : geolinguistic profiles and scenarios of language contact in India. Presses de l'Université Laval; 1991.

52. Segato R. Discurso inaugural de Rita Segato - YouTube. Feria Internacional del Libro de Buenos Aires; 2019.

53. Shashok K. Editing around the World AuthorAID in the Eastern Mediterranean: a communication bridge between mainstream and emerging research communities. European Science Editing. 2009.

#### Información suplementaria

**S1 Artículo completo en español.**

**S2 Resumen de los resultados obtenidos organizados por su impacto en investigadores colombianos en español.** Segundo idioma o idioma extranjero (L2), primer idioma o lengua materna (L1).

**S3 Preguntas de encuesta.** Preguntas en español de la encuesta "Implicaciones del lenguaje en publicaciones científicas".
