## Supplementary figures and images for "Disadvantages of writing, reading, publishing and presenting scientific papers caused by the dominance of the English language in science: The case of Colombian PhD in biological sciences"

### S2 Summary of the results obtained organized by their impact on Colombia researchers in Spanish.

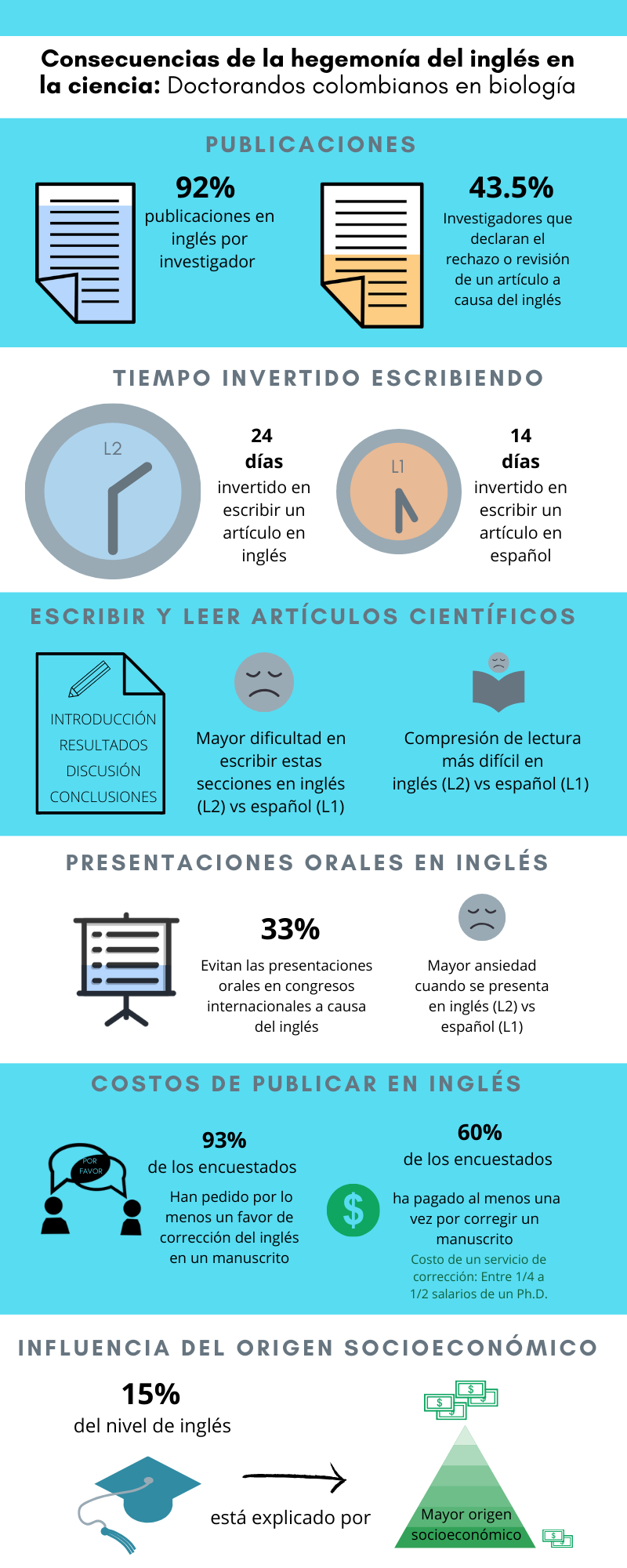
