## Supplementary material for "Disadvantages of writing, reading, publishing and presenting scientific papers caused by the dominance of the English language in science: The case of Colombian PhD in biological sciences": S3 Survey questions.

### Implicaciones del idioma en las publicaciones científicas

Queridos(as) colegas,

Con esta encuesta están contribuyendo de manera anónima a una tesis de la Maestría en Comunicación Científica de la Universidad de Barcelona. El proyecto es sobre la importancia del multilingüismo en las publicaciones científicas, especialmente en el proceso de escritura de artículos.

Esta encuesta está dirigida a personas hispanohablantes (con enfásis en colombianos(as)) que estén realizando o hayan finalizado un doctorado en ciencias naturales y básicas (física, química, biología, biomedicina, etc).

Agradezco fuertemente su participación y su posterior difusión a otros(as) colegas.

*Obligatorio

#### Datos básicos

1. **Nacionalidad** *

2. **País en el que vive**
No ciudad o municipio

3. **Edad**

4. **Tipo de institución donde realizó sus estudios de bachillerato o secundaria**
*Marca solo un óvalo.*
- Colegio público
- Colegio privado

5. **Tipo de institución donde realizó sus estudios universitarios de grado**
*Marca solo un óvalo.*
- Universidad pública
- Universidad privada

6. **Escoja la ocupación a la que pertenece:**
*Marca solo un óvalo por fila.*

| | Grandes empleadores, los altos directivos de las empresas y la Administración Pública, profesionales de alto nivel | Directivos y profesionales de nivel bajo | Empleados de nivel alto | Pequeños empleadores y por los trabajadores autónomos no profesionales | Trabajadores autónomos agrícolas | Supervisores y técnicos de rango inferior | Trabajadores del comercio y los servicios de rango inferior | Trabajadores manuales cualificados | Trabajadores no cualificados | Excluidos c mercado laboral, desemplead sin experienci previa |
|---|---|---|---|---|---|---|---|---|---|---|
| Madre | | | | | | | | | | |
| Padre | | | | | | | | | | |
| Abuelos(as) maternos | | | | | | | | | | |
| Abuelos(as) paternos | | | | | | | | | | |
| Hermano(a) 1 | | | | | | | | | | |
| Hermano(a) 2 | | | | | | | | | | |
| Hermano(a) 3 | | | | | | | | | | |

7. **Estrato social de la vivienda (Solo si viven en Colombia)**
*Marca solo un óvalo por fila.*

| | 1 | 2 | 3 | 4 | 5 | 6 |
|---|---|---|---|---|---|---|
| Usted | | | | | | |
| Madre | | | | | | |
| Padre | | | | | | |
| Abuelos(as) paternos | | | | | | |
| Abuelos(as) maternos | | | | | | |
| Hermano(a) 1 | | | | | | |
| Hermano(a) 2 | | | | | | |
| Hermano(a) 3 | | | | | | |

8. **Primera lengua(s)** *
En caso de ser bilingüe o trilingüe escriba 2 idiomas que principalmente usa

9. **Idioma(s) que domina**

10. **Nombre del programa o campo del doctorado que realiza o realizó** *

11. **Universidad o instituto dónde realiza o realizó el doctorado**

12. **Ya finalizó el doctorado?**
*Marca solo un óvalo.*
- Sí
- No

13. **Si ya finalizó el doctorado, cuál es su ocupación actual?**

14. **¿En qué año inició el doctorado?**

15. **Número aproximado de publicaciones académicas que tiene** *
Como autor principal y coautor. Artículos publicados, en revisión o aceptadas

16. **Porcentaje de sus publicaciones que ha realizado en INGLÉS** *
Como autor principal y coautor. Artículos publicados, en revisión o aceptadas
*Marca solo un óvalo.*
- Ninguna
- 10%
- 20%
- 30%
- 40%
- 50%
- 60%
- 70%
- 80%
- 90%
- 100%

17. **Porcentaje de sus publicaciones que ha publicado en su PRIMERA LENGUA**
Como autor principal y coautor. Artículos publicados, en revisión o aceptadas
*Marca solo un óvalo.*
- Ninguna
- 10%
- 20%
- 30%
- 40%
- 50%
- 60%
- 70%
- 80%
- 90%
- 100%

#### Escritura de artículos en inglés

18. **Una vez ya tiene completos los resultados de un estudio, cuánto tiempo se demora en el proceso de escritura del artículo si este se va a publicar en INGLÉS?**
Tiempo en días y número de horas diarias dedicadas. Ej: 20 días, 2 horas diarias

19. **Una vez ya tiene completos los resultados de un estudio, cuánto tiempo se demora en el proceso de escritura del artículo si este se va a publicar en su PRIMERA LENGUA?**
Tiempo en días y número de horas diarias dedicadas. Ej: 20 días, 2 horas diarias

20. **Al escribir un artículo que se va a publicar en inglés, usted preferentemente?** *
*Marca solo un óvalo.*
- Escribe en inglés directamente
- Escribe en su primera lengua y posteriormente hace la traducción al inglés

21. **Ha pedido favores o pagado por CORRECIÓN o REVISIÓN del inglés un artículo?** *
Sólo CORRECCIONES y EDICIÓN de texto, no traducciones
*Marca solo un óvalo.*
- Sí, he pedido favores
- Sí, he pagado
- Sí, ambas posibilidades
- No

22. **¿Cuál es el porcentaje de artículos por los cuales ha pedido favores o/y pagado por CORRECIÓN o REVISIÓN del inglés?**
Sólo CORRECCIONES y EDICIÓN de texto, no traducciones
*Marca solo un óvalo por fila.*

| | Ninguno | 10% | 20% | 30% | 40% | 50% | 60% | 70% | 80% | 100% |
|---|---|---|---|---|---|---|---|---|---|---|
| Pedido un favor | | | | | | | | | | |
| Pagado | | | | | | | | | | |

23. **Si ha pagado, aproximadamente cuánto paga por cada artículo?**
Sólo CORRECCIONES y EDICIÓN de texto, no traducciones. En dólares

24. **Si ha pagado, cuál es la procedencia de ese dinero?**
Sólo CORRECCIONES y EDICIÓN de texto, no traducciones
*Marca solo un óvalo.*
- Únicamente yo
- Yo y los coautores
- Institución(es) académica(s)
- Beca(s) o subvención(es)
- Recursos mixtos

25. **Ha pedido favores o pagado por TRADUCIR un artículo al inglés?** *
Sólo traducciones
*Marca solo un óvalo.*
- Sí, he pedido favores
- Sí, he pagado
- Sí, ambas posibilidades
- No

26. **¿Cuál es el porcentaje de artículos por los cuales ha pedido favores o/y pagado por la TRADUCCIÓN al inglés?**
Sólo traducciones
*Marca solo un óvalo por fila.*

| | Ninguno | 10% | 20% | 30% | 40% | 50% | 60% | 70% | 80% | 100% |
|---|---|---|---|---|---|---|---|---|---|---|
| Pedido un favor | | | | | | | | | | |
| Pagado | | | | | | | | | | |

27. **Si ha pagado, aproximadamente cuánto paga por cada artículo?**
Sólo TRADUCCIONES. En dólares

28. **Si ha pagado, cuál es la procedencia de ese dinero?**
Sólo TRADUCCIONES
*Marca solo un óvalo.*

- Únicamente yo
- Yo y los coautores
- Institución(es) académica(s)
- Beca(s) o subvención(es)
- Recursos mixtos

29. **Alguna vez ha dejado de participar (o de enviar el resumen para participar) en un congreso internacional por el hecho que debía hablar en inglés?**
*Marca solo un óvalo.*

- Sí
- No
- No quise una hacer presentación oral
- No quise presentar póster

30. **Califique que tan ansioso se siente por hablar en un congreso?**
*Marca solo un óvalo por fila.*

| | Nada | Poco | Regular | Muy | Extremadamente |
|---|---|---|---|---|---|
| Presentando en inglés | | | | | |
| Presentado en su primera lengua | | | | | |

31. **En el momento de leer un artículo científico en INGLÉS usted cree que ...**
*Marca solo un óvalo por fila.*

| | Muy difícil | Difícil | Regular | Fácil | Muy fácil |
|---|---|---|---|---|---|
| Entender el texto en general | | | | | |
| Entender la terminología científica es | | | | | |
| Interpretar figuras y tablas es | | | | | |

32. **En el momento de leer un artículo científico en PRIMERA LENGUA usted cree que ...**
*Marca solo un óvalo por fila.*

| | Muy difícil | Difícil | Regular | Fácil | Muy fácil |
|---|---|---|---|---|---|
| Entender el texto en general | | | | | |
| Entender la terminología científica es | | | | | |
| Interpretar figuras y tablas es | | | | | |

33. **En el momento de escribir un artículo científico en INGLÉS usted cree que ...**
*Marca solo un óvalo por fila.*

| | Muy difícil | Difícil | Regular | Fácil | Muy fácil |
|---|---|---|---|---|---|
| Escribir la introducción es | | | | | |
| Escribir la metodología es | | | | | |
| Escribir los resultados es | | | | | |
| Escribir la discusión es | | | | | |
| Escribir la conclusión es | | | | | |

34. **En el momento de escribir un artículo científico en su PRIMERA LENGUA usted cree que ...**
*Marca solo un óvalo por fila.*

| | Muy difícil | Difícil | Regular | Fácil | Muy fácil |
|---|---|---|---|---|---|
| Escribir la introducción es | | | | | |
| Escribir la metodología es | | | | | |
| Escribir los resultados es | | | | | |
| Escribir la discusión es | | | | | |
| Escribir la conclusión es | | | | | |

35. **Prefiere leer textos científicos en su primera lengua**
*Marca solo un óvalo.*

- Sí / Yes
- No

36. **Alguna revista le ha rechazado o solitado la correción de un artículo científico por una escritura incorrecta del inglés** *
*Marca solo un óvalo.*

- Sí / Yes
- No

37. **Sí tiene algún ejemplo de comentarios de los editores con respecto a la escritura incorrecta del inglés, por favor escríbalo**

#### Aprendizaje de inglés

38. **Qué porcentaje de inglés habla diariamente?**
*Marca solo un óvalo.*

- No hablo inglés diariamente
- 10%
- 20%
- 30%
- 40%
- 50%
- 60%
- 70%
- 80%
- 90%
- 100%

39. **Desde qué edad comenzó a aprender inglés?** *

40. **Cuántos años estudió o lleva estudiando inglés?** *

41. **Califique su experiencia de aprendizaje en la institución(es) donde estudió inglés** *
*Marca solo un óvalo.*

| | 1 | 2 | 3 | 4 | 5 | |
|---|---|---|---|---|---|---|
| Mala | | | | | | Muy buena |

42. **Cuántos años ha vivido en un país angloparlante o que utilizará el inglés como primer idioma?**
Si nunca ha vivido coloque cero

43. **Califique su nivel de inglés actual** *
*Marca solo un óvalo por fila.*

| | Excelente | Bueno | Regular | Malo |
|---|---|---|---|---|
| Escritura | | | | |
| Habla | | | | |
| Escucha | | | | |
| Lectura | | | | |

44. **Ha presentado alguna examen de inglés con reconocimiento internacional?**
*Marca solo un óvalo.*

- Sí
- No

Con la tecnología de Google Forms
